# Effects of larval diet on the life-history traits and phenotypic expression of pyrethroid resistance in the major malaria vector *Anopheles gambiae s.s.*

**DOI:** 10.1101/2022.01.11.475801

**Authors:** Pierre Marie Sovegnon, Marie Joelle Fanou, Romaric Akoton, Oswald Yédjinnavênan Djihinto, Hamirath Odée Lagnika, Romuald Agonhossou, Luc Salako Djogbénou

**Affiliations:** Tropical Infectious Diseases Research Centre, University of Abomey-Calavi, Cotonou, Benin; Regional Institute of Public Health/University of Abomey-Calavi, Benin; Department of Vector Biology, Liverpool School of Tropical Medicine, Pembroke Place, Liverpool L3 5QA, United Kingdom

**Keywords:** Larval diet, Life-history traits, Pyrethroid resistance, *Anopheles gambiae s.s*.

## Abstract

The success achieved in reducing malaria transmission by vector control is threatened by insecticide resistance. To strengthen the current vector control programmes, the non-genetic factors underlying the emergence of insecticide resistance in *Anopheles* vectors and its widespread need to be explored. This study aimed to assess the effects of larval diet on some life-history traits and pyrethroid-insecticide susceptibility of *Anopheles gambiae s.s*.

Three (3) *An. gambiae* strains, namely Kisumu (insecticide susceptible), AcerKis (homozygous *ace-1*^R^G119S resistant) and KisKdr (homozygous *kdr*^R^L1014F resistant) were fed with three different diets (low, medium, and high) of TetraMin^®^ Baby fish food. Pre-imaginal developmental time, larval mortality, adult emergence rate and female wing length were measured. Mosquito females were exposed to insecticide-treated net (ITN) PermaNet 2.0 and PermaNet 3.0.

In the three *An. gambiae* strains, significant differences in adult emergence rates (*F* = 1054.2; *df* = 2; *p* <0.01), mosquito wing length (*F* = 970.5; *df* = 2; *p* <0.01) and adult survival post insecticide exposure (*χ^2^*= 173; *df* = 2; *p* <0.01), were noticed among the three larval diets. Larvae fed with the low food diets took more time to develop, were smaller at emergence and displayed a short lifespan, while the specimens fed with a high regime developed faster and into big adults. Although being fed with a high diet, none of *An. gambiae* strain harbouring the *kdr*^R^ and *ace-1^R^* allele survived 24 hours after exposure against PermaNet 3.0.

This study showed that variation in the larval diet significantly impacts *An. gambiae* life-history traits such as larval mortality and developmental time, adult wing length, and female susceptibility to pyrethroid insecticides. Further investigations through field-based studies would allow an in-depth understanding of the implications of these non-genetic parameters on the physiological traits of malaria vectors and consequently improve resistance management.

## Introduction

Malaria is a parasitic disease that is still raising public health concerns, and more than 90% of malaria cases still occur in Africa [1]. The causative agent of this disease is transmitted by infected *Anopheles* female mosquitoes, which carry the infection to humans [2]. In Sub-Saharan Africa, *An. gambiae*, *An. arabiensis*, *An. coluzzii* and *An. funestus* are the dominant malaria vectors [3,4].

Among the malaria control strategies, vector control remains the primary component [5], through the widespread use of Long-lasting insecticide-treated nets (LLINs) and Indoor Residual Spraying (IRS) [6,7]. These interventions reduced human-vector contact and, consequently, malaria transmission [8]. However, the phenomenon hampering the effectiveness of these core interventions is the emergence and spread of insecticide resistance mechanisms in malaria vector populations. Resistance to pyrethroid insecticides mainly used for LLINs has been reported in *An. gambiae* mosquitoes across several African countries [9–11] and appears to rely mainly on metabolic resistance mechanisms and target-site modification [12,13]. The most described target modifications are the mutations in the voltage-gated sodium channel gene (L1014F, L1014S and N1575Y) which confer resistance to pyrethroids and Dichloro-Diphenyl-Trichloroethane (DDT) [14–16] and mutation in the *Ace-1* gene (G119S) conferring resistance to organophosphates and carbamates [17]. These hinder the effectiveness of the current tools used in malaria vector control.

Resistance phenotype is an interaction between environmental factors and genotype determining the resistance profile [18]. In organisms with complex life cycles such as mosquitoes, environmental factors affecting earlier life stages could impact the following stages [18,19]. Several studies have revealed that larval environment determines the adult morphological and physiological features such as adult size, biting behaviour, fecundity, longevity, and vector competence [20–22]. Availability of food is one of these environmental factors that might influence mosquito life-history traits [23]. It was reported that larval nutritional conditions could have carry-over effects on adult life-history traits and the susceptibility to other abiotic stresses such as insecticides [24]. Indeed, it was reported that *Anopheles coluzzii* larval developmental time was longer at lower food concentrations and the emerged adults were much smaller [25]. Furthermore, whereas it was shown that the expression of metabolic resistance mechanism against DDT was not affected by larval food deprivation [26], *Anopheles arabiensis* mosquitoes carrying L1014S *kdr* allele emerged from underfed larvae were less tolerant to DDT [27]. Therefore, the real pattern of insecticide resistance profile in malaria vectors might be influenced by larval nutritional conditions leading to biased decisions when designing malaria vector control strategies.

However, in *An. gambiae* carrying target-site insensitivity *kdr^R^* (L1014F) and *ace-1^R^* (G119S) alleles, no study has been carried out to determine whether and how the dietary resources at the pre-imaginal stage might influence the pyrethroid-resistance phenotypes. Moreover, the influence of larval food resources on the different stages of larval development in these resistant mosquitoes remain unknown. The present study aimed to assess the effect of different larval diets on life-history traits and phenotypic expression of pyrethroid resistance in *An. gambiae* mosquitoes. Such information would greatly benefit our understanding of the evolution of resistance and could advise strategies for vector control initiatives.

## Material and methods

### Mosquito strains

Three *An. gambiae* laboratory strains were used in this study. Kisumu, sampled from Kenya in the early 1950s, is a reference strain susceptible to all insecticides [28]. AcerKis is homozygous (*Ace-1^RR^*) for the G119S allele and resistant to organophosphate and carbamate insecticides [29]. KisKdr strain which is homozygous (*kdr^RR^*) for the L1014F allele and conferring resistance to pyrethroids and DDT and was obtained by introgression of the *kdr^R^* (L1014F) allele into the Kisumu genome [30]. Both AcerKis and KisKdr were supposed to share the same genetic background as the Kisumu strain and were free of metabolic resistance [29].

### Mosquito rearing and larval diet

The three *An. gambiae* strains were reared in the Laboratory of Infectious Vector-Borne Diseases (LIVBD) of Regional Institute of Public Health (IRSP, Benin), under standard insectary conditions (insecticide-free environment, 25-27°C room temperature, 70-80% relative humidity and 12:12 light and dark period). They were fed daily by TetraMin^®^ Baby fish food.

Each mosquito strain’s first instar larvae (L1) were initially randomly selected and assigned to three experimental groups based on the amounts of food provided, according to Kulma et al. [26]. The first group was reared under plentiful food conditions (hereafter, “high-diet”), the second group was reared under standard food conditions (hereafter, “medium-diet”), and the third group was reared under scare food conditions (hereafter, “low-diet”).

A total of 900 tubes (37.5 mm in diameter and 65 mm in height), each containing three (3) first instar larvae (L1), were used over three replicates per mosquito strain. Each replicate was set up as follow: 300 tubes (100 for low-diet, 100 for medium-diet and 100 for high-diet). Each tube contained 10 mL of dechlorinated water, and food was provided daily at the same hour. To partially offset water evaporation, the food was diluted in 100 μL of dechlorinated water. Water was changed every two (2) days until larvae reached the fourth instar stage to avoid uncontrolled food accumulation. At the pupation stage, pupae were transferred to an emergence cage (30×30×30 cm), and the resulting adults were counted and provided with a 10% honey solution on cotton wool pads.

### Larval mortality and development time

To determine the larval mortality rate associated with the larval diet, dead larvae were counted daily in each food condition during the larval cycle. The timing of adult emergence was also recorded, and the larval development time was estimated as the period elapsed from the L1 stage to emergence.

### Adult bioassay

Susceptibility to the insecticide-treated nets of the emerged adult females was assessed by performing the WHO cone test. The cone test is an adaptation of the WHO cone bioassay, with the following modification: During the assay, the test operator holds his forearm behind the cone (**Fig 1**) to attract mosquitoes towards the pieces of the net as described by Bohounton et al. [31].

**Fig 1.**
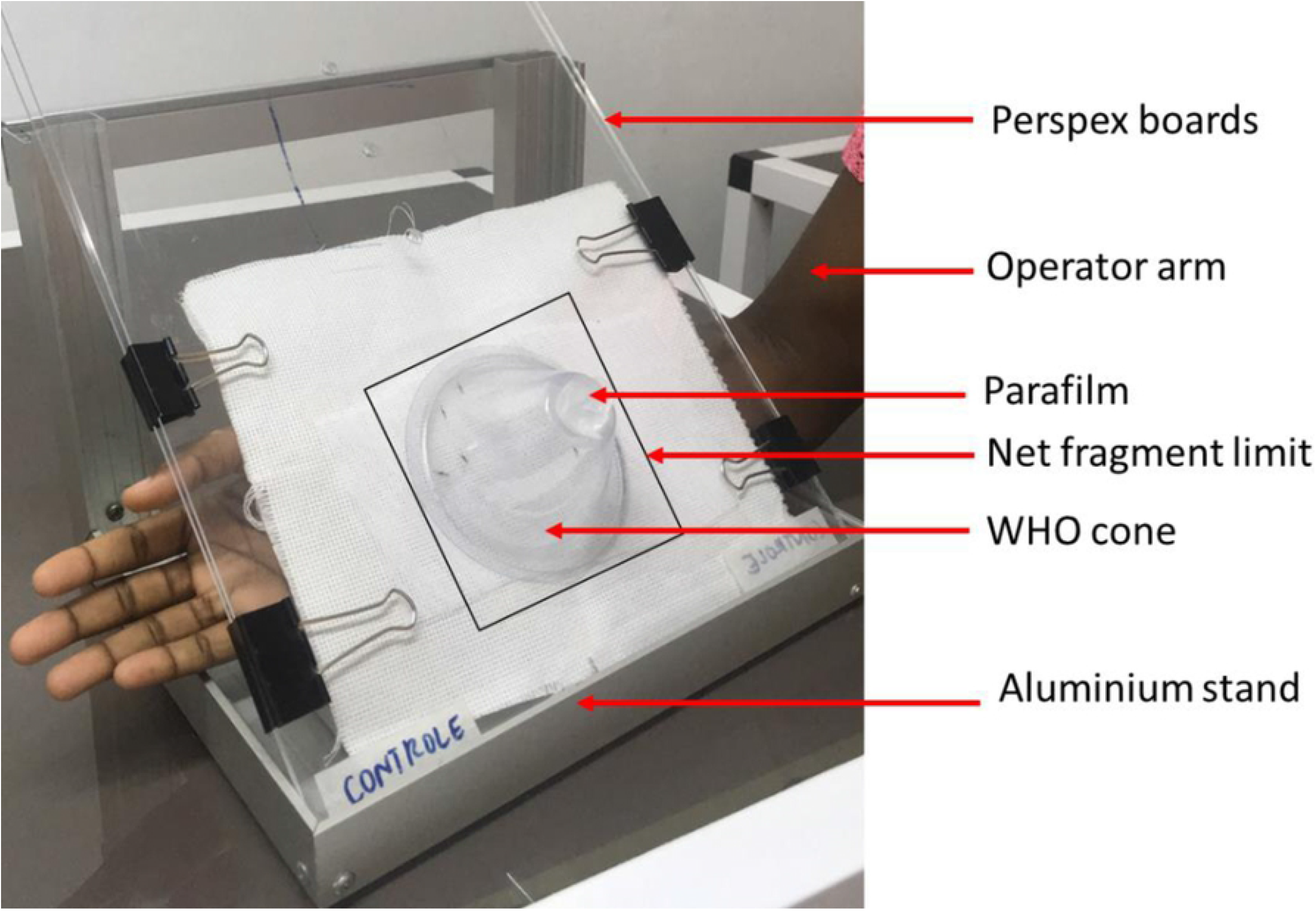
Cone test equipment. (Image reproduced from Bohounton et al. [30] with permission of the corresponding author)

Nets pieces (13 cm x 13 cm; 169 cm^2^) were prepared for each of the following insecticide-treated nets: PermaNet 2.0 (net treated at 55 mg/m^2^ of deltamethrin) (Vestergaard, Lausanne, Switzerland) and PermaNet 3.0 (side panel treated at 85 mg/m^2^ of deltamethrin) (Vestergaard, Lausanne, Switzerland). Nets of the same size without insecticide treatment (Coghlan’s) were also used as a control. A batch of 5 mosquitoes was transferred into the cone with the operator’s forearm in position. Mosquitoes were then exposed to the insecticide-treated and control net pieces for three minutes [31].

After exposure, mosquitoes were removed from the cone, transferred into a recovery cup, and provided with 10% of honey solution soaked on a cotton pad. Mosquito mortality was then recorded every day until the death of the last female of each mosquito strain.

### Measurement of wing length

Wing length was used to measure mosquitoes’ body size [32]. After female mosquitoes’ exposure to the insecticide-treated nets, the wings of dead mosquitoes were removed, and the length of each wing was measured from the tip to the distal end of the alula (excluding the fringe) [32] using Image *Hotviewer* software (version 2.0).

### Ethical statement

This study was approved by the Research Ethics Committee, University of Abomey-Calavi, Benin (N^O^: 126 of 12/02/2020)

### Data analysis

All data generated were recorded in Microsoft Excel sheets and exported to R 3.5 [33] and GraphPad Prism 8.0.2 software (San Diego, California USA) for analysis. The normality of data distribution was checked using the Shapiro Wilk test [34]. To assess the influence of main variables (strains and food conditions) and their interaction on the larval mortality rates, the emergence rates and the wings length, a two-way analysis of variance test (ANOVA) was used, followed by Tukey multiple pairwise-comparisons test. The Larval developmental time was analysed using a Generalised Linear Mixed Model (GLMM) with a binomial error distribution. The mosquito strain, food diet conditions, and interaction were coded as fixed factors.

The effect of diets on the resistance phenotype of *An. gambiae s.s*. was assessed by comparing mosquito survival in each diet regime. Kaplan-Meier survival curves analysed the survival of adult mosquitoes after insecticide exposure. The log-rank test was performed to assess the difference in survival between the three strains. The effect of the nets (PermaNet 2.0 and PermaNet 3.0) and the food diet conditions on the longevity were analysed using Cox’s proportional hazard models. All statistical analyses were set at a significance threshold of *p* < 0.01.

## Results

### Larval mortality rate

Significant influence of the diet conditions on the larval mortality rate of the mosquito strains (*F* = 1113.75; *df* = 2; *p* < 0.01) was displayed. An overall increased larval mortality was observed in each of the three mosquito strains fed with the lowest amount of food (**Fig 2A**). The highest larval mortality rate was recorded in AcerKis larvae (83.58%; 81–0.85), which is significantly different from that of Kisumu (70.5%; 95% CI: 0.68-0.73) and KisKdr (72.16%; 95% CI: 0.67–0.74). However, no difference in the mortality rate was observed between Kisumu and KisKdr (*F* = 0.037; *df* = 1; *p* = 0.848). With the high larval diet, the only difference in mortality rates was observed between Kisumu and AcerKis larvae (*F* = 19.05; *df* = 1; *p* < 0.01).

**Fig 2.**
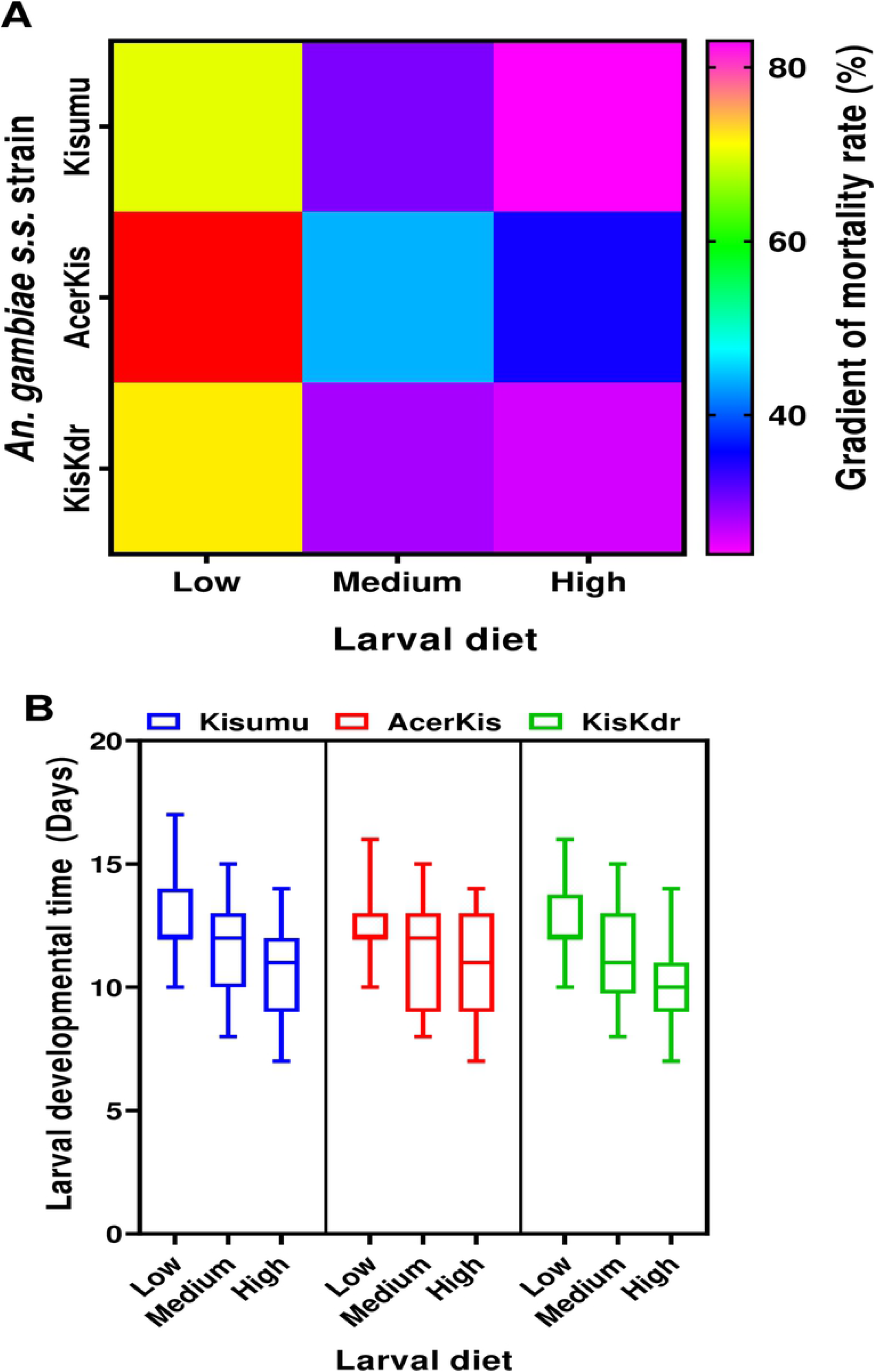
Effect of larval diets on the mortality and developmental time of *Anopheles gambiae s.s*. larvae. **A:** Larval mortality rates related to larval diets. The mortality rates are illustrated by a multi-colour gradient from purple (0%) to red (83% maximum). **B:** Larval developmental time for each mosquito strain and larval-feeding regime. Thick horizontal lines represent the median, bottom and upper edges of the boxes are first and third quartiles, whiskers indicate minimum and maximum values.

### Larval developmental time

Overall, the length of larval developmental time varies between 10 and 17 days for all mosquito strains and nutritional regimes. Significant influence of the diet conditions on the larval developmental time rate of the mosquito strains (*F* = 435.67; *df* = 2; *p* < 0.01) was also displayed. The larvae fed with the high food diet developed significantly faster than did their counterparts fed with the low food regime within each mosquito strain **(Fig 2B)**.

### Adult emergence rate

The adult emergence rate was significantly affected by the nutritional regime within all the three *An. gambiae* s.s. strains (*F* = 1054.206; *df* = 2; *p* < 0.01). In all mosquito strains analysed, very few adults emerged from larvae fed with the low diet **(Fig 3A)**, with the lowest emergence rate in AcerKis larvae (14.83 ± 2.01%) compared to Kisumu (25.33 ± 2.5%) and KisKdr larvae (24.83 ± 2.5%). Inversely, significantly high proportions of adults were recorded from larvae fed with the high food diet (66.91 ± 2.6% in AcerKis; 67.58 ± 2.6% in KisKdr and 64.16 ± 2.7% in Kisumu).

**Fig 3.**
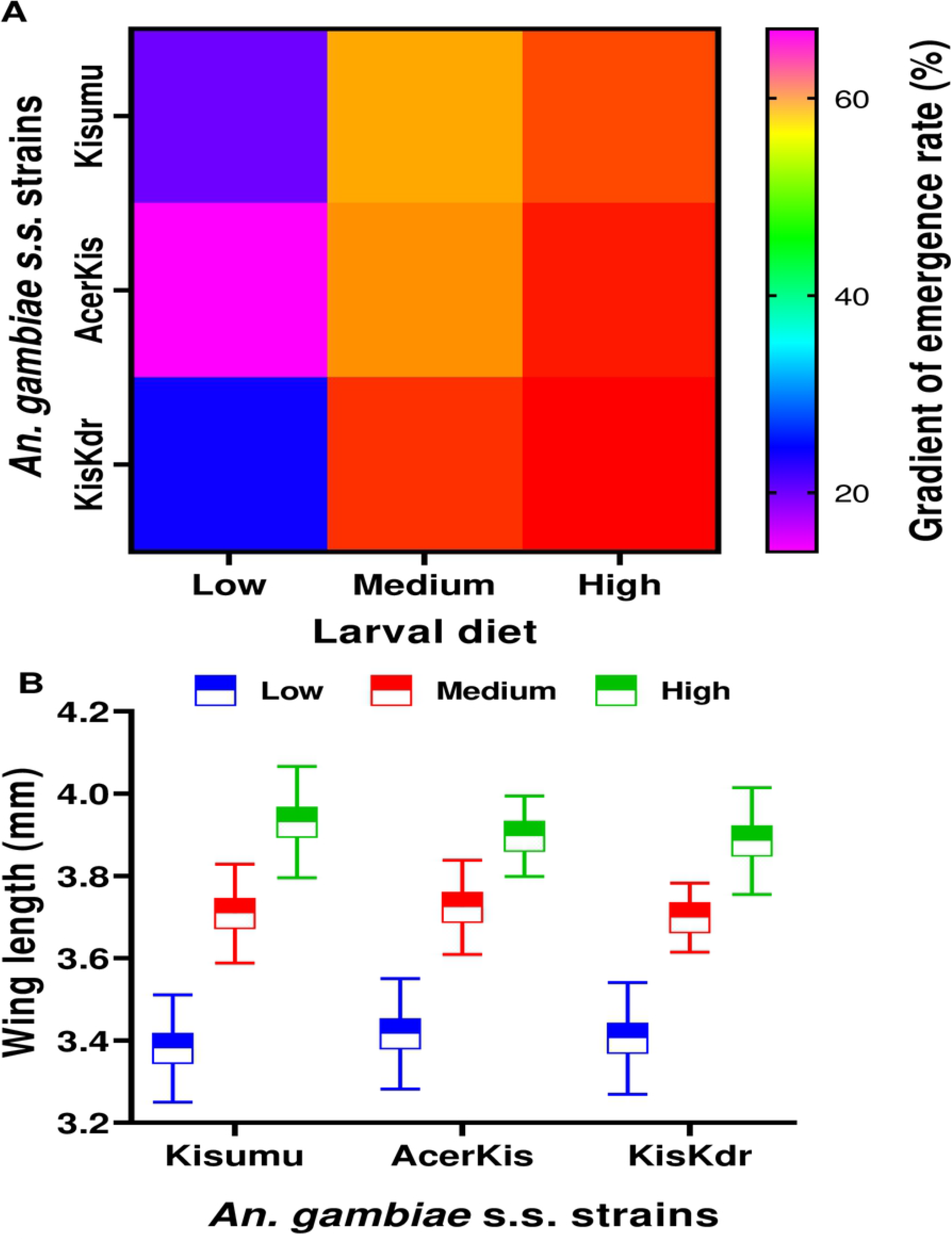
Influence of larval diets on emergence rate and body size in *Anopheles gambiae* s.s. **A:** Adult emergence rate for each mosquito strain and feeding regime. The mortality rates are illustrated by a multi-colour gradient from purple (0%) to red (67% maximum). **B:** Wing length of adult females emerged from each larval-diet treatment.

### Adult mosquitoes wing length

There was a significant variation of wing length according to the diet (*F* = 970.571; *df* = 2; *p* <0.01). A significantly higher wing length was observed in adults that emerged from larvae fed on the high amount of food (4.0 ± 3.8 mm) compared to those fed on medium (3.8 ± 3.6 mm) and low amount (3.4 ± 3.2 mm) of food **(Fig 3B)**.

### Adult mosquitoes’ survivorship post-exposure to the insecticide-treated nets

Lower adult emergence rate was recorded in the low diet larvae categories; consequently, there were few adult females for bioassay. Thus, only females from medium and high food regimes were used for insecticide susceptibility tests.

Kaplan-Meier survival curves analysis showed that in each mosquito strain, adult females from the medium diet had a shorter lifespan and median survival time than females fed on the high diet in the absence of insecticide (untreated control net) **(Fig 4)**.

**Fig 4.**
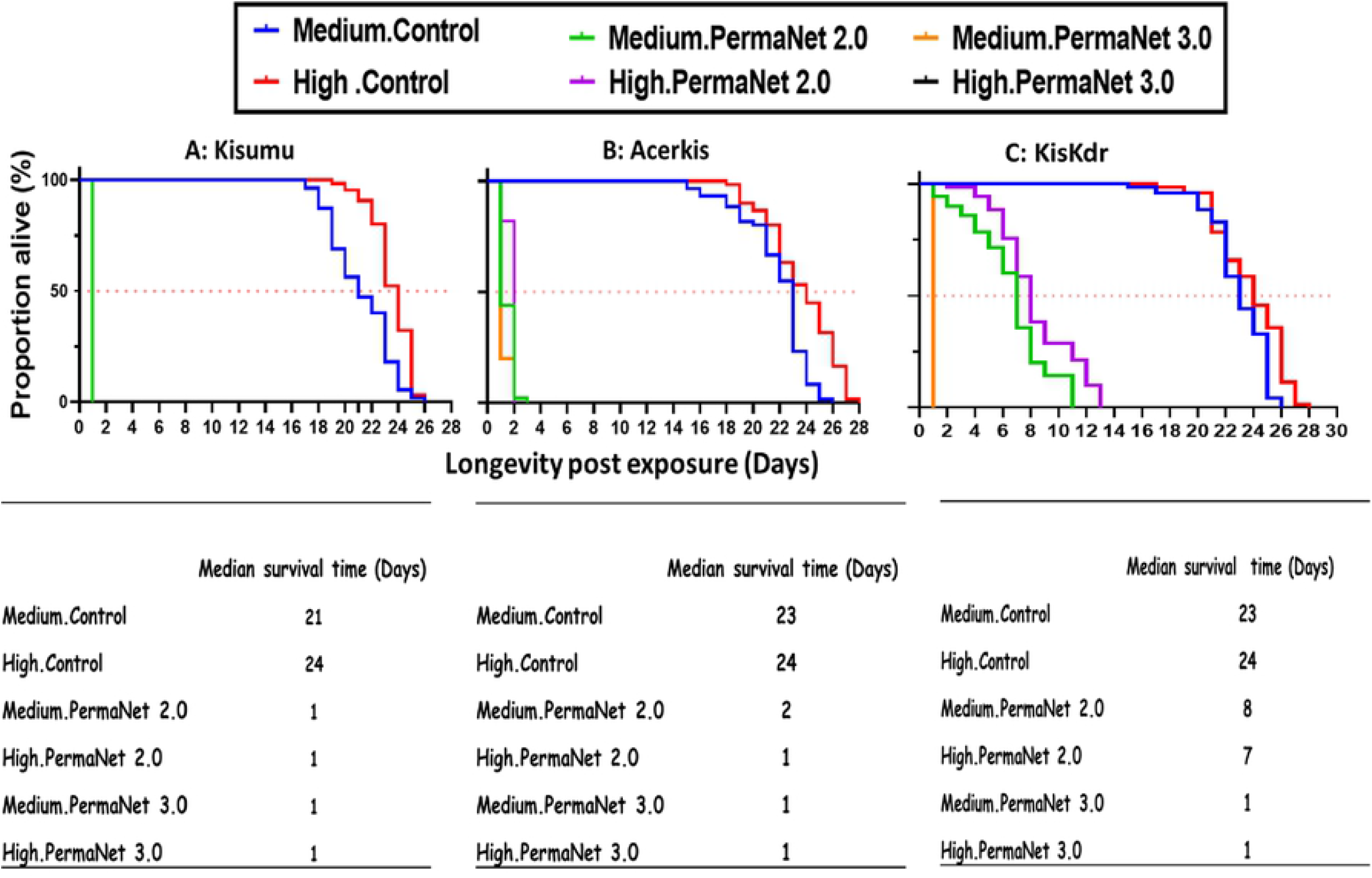
Mosquito survivorship post-exposition to insecticide treated nets. The Kaplan Meier survival curves for each mosquito strain from medium and high food regimes exposed to the different insecticide treated net pieces are represented in **(A)** for Kisumu, **(B)** for AcerKis, and **(C)** for KisKdr. The median survival times of each mosquito strain in each experimental condition were shown under the corresponding panel.

In the Kisumu strain, no adult females survived 24 hours after exposure to PermaNet 2.0 and PermaNet 3.0 **(Fig 4A).** A significant decrease in survivorship of AcerKis females from medium and high diets compared to the control (*χ^2^* = 173; *df* = 2; *χ^2^* = 173; *p* < 0.01). AcerKis females from the medium diet exposed to PermaNet 2.0 had lived up to three days and two days when exposed to PermaNet 3.0 with the same median survival time of 1 day **(Fig 4B)**. For those from the high diet exposed to the PermaNet 2.0 and PermaNet 3.0, the survival of mosquitoes was two days with a median survival time of 1 day.

KisKdr females from high diet displayed significantly reduced survival when exposed to PermaNet 2.0 (13 days) compared to their control with a median survival time of 8 days **(Fig 4C)**. This survivorship decreased significantly to 11 days with a median survival time of 7 days in females from the medium diet regime (*χ^2^* = 7.2; *df* = 1; *p* < 0.01). However, all females from medium and high diets exposed to PermaNet 3.0 died within 24 hours.

## Discussion

The pre-imaginal period is a critical stage of the malaria-transmitting mosquitoes life cycle [35]. Thus, it appears critical to improve our understanding concerning the mosquito larval bio-ecology for the effectiveness of vector control measures in the endemic areas. Indeed, the environment in which an individual develops itself can considerably impact its adult phenotype through carry-over effects. This is of obvious relevance in vector-borne diseases such as malaria since any changes in mosquito life-history traits could have significant involvements for the parasite transmission through changes in key parameters of vectorial capacity [22,36]. The present study investigated whether larval nutrition can influence life-history traits and the expression of pyrethroid resistance in *An. gambiae* mosquito harbouring only target sites insensitivity *kdr^R^* and *Ace-1^R^* alleles.

In this study, among life-history traits such as larval development, mosquito density, female size and pyrethroid insecticide tolerance were shown to be affected by larval diet. Indeed, we observed that, when fed with the low food diet, larvae of all three mosquito strains used took more time to develop, with increased mortality rates when compared to those fed with the high diet. Consequently, the high mortality rates in low nutritional conditions led to low emergence rates in all mosquito strains. Strategies towards food deprivation in the larval environment that could lead to decreased vector density, may benefit the malaria vector control. Furthermore, in the field, a longer larval developmental time observed with the low food diet could expose larvae to other stressors or risks factors such as competition [37], predation [38] or drought [39], which are natural factors that would contribute to the decreasing of larval density in breeding sites. The inability to develop until pupation and the low emergence rate could be associated with a low reserve accumulation due to the low amount of nutrients, as observed in this study. In addition, the highest mortality rates were recorded in the resistant AcerKis larvae fed with low diet and high larval-food conditions when compared to other strains. Also, even with the high food regime, their developmental time was longer than in the resistant strain KisKdr. These observations suggest that the AcerKis larvae need to be more fed and consequently could need more nutrients for their optimal development compared to other strains, and even so, the high amounts of food used in this study were not yet sufficient to ensure rapid larval development. This sensitivity to the diet variations might be due to an additional genetic cost linked to the *Ace-1^R^* allele [40].

This study also showed that larval diets were positively correlated to the adult mosquito size. Adult mosquitoes from low food diets were much smaller than those from high food conditions. Previous research reported that dietary resources in larval habitats determine the size of adult mosquitoes and most often account for the differences in developmental fitness of the emerged adult [42]. The low food regimes used in this study likely contain a meagre amount of nutrients that contributed to the low reserves accumulation of protein, lipid, and glycogen, resulting in small-sized teneral adults. It was shown that tiny female mosquitoes need two or three blood meals to complete the first gonotrophic cycle [43, 44]. Also, smaller mosquitoes produce fewer eggs during their lifetime than their larger counterparts [46].

The effect of the different larval diet regimes on adult mosquitoes’ insecticide susceptibility was also assessed. The amounts of food available at the larval stage influenced the expression of mosquitoes’ resistance to deltamethrin, the main insecticide coated on PermaNet 2.0 and PermaNet 3.0 bednets. KisKdr females from high diet conditions survived more than KisKdr females from medium diet regimes when exposed to PermaNet 2.0. They probably would accumulate more energy reserves that allowed them to tolerate more doses of insecticides. The previous study has shown no variability in the phenotypic expression of insecticide resistance according to the larval diets in the strain of *An. gambiae s.s*. ZAN/U strain is characterised by enzymatic detoxification resistance [26]. Therefore, resistance by enzymatic resistance mechanisms would be less sensitive to variations in larval diet and would require fewer energy resources than resistance mechanisms by mutations in the *Kdr* gene [26].

A long survival time was recorded in mosquitoes fed with the high food diet in deltamethrin-free conditions. Female lifespan is a significant epidemiological factor as it considers vectorial capacity [20]. Only infected mosquitoes that survive long enough can transmit the *Plasmodium* parasites responsible for the disease onset [41]. Consequently, the prolonged survival time observed with the high food diet in the three mosquito strains in the absence of insecticide seems to be a contrasting epidemiological output. Even so, the fecundity, fertility, and *Plasmodium* infection susceptibility of these mosquitoes were not evaluated in the current study and deserved further consideration.

Furthermore, the susceptible (Kisumu) and resistant (AcerKis, KisKdr) strains survival decreased significantly when exposed to both PermaNet 2.0 and PermaNet 3.0 bednets (when compared to the control nets), suggesting that although being well-fed, *An. gambiae* strains harbouring *Kdr^R^* and *Ace-1^R^* alleles were less tolerant to the insecticide-treated bednets. KisKdr females from high diet conditions survived significantly longer when exposed to PermaNet 2.0 than those exposed to the PermaNet 3.0 bednets. The latter is not a surprise since it was already demonstrated that PermaNet 2.0 had shown less efficacy against resistant *Anopheles* mosquitoes [42,43]. In addition, the next-generation net, PermaNet 3.0, was found to be more efficient against pyrethroid-resistant *Anopheles* mosquitoes [44].

## Conclusion

This study showed that larval nutritional stress affects the life-history parameters measured under laboratory conditions in *Anopheles gambiae*. However, although being well-fed, *An. gambiae* strains harbouring *kdr*^R^ and *ace-1^R^* alleles were less tolerant to the pyrethroid insecticide. Given the findings of this study, larval nutritional stresses could be implemented as a vector control tool. The genotype-environment interaction needs to be further investigated to understand the evolution of resistance and provide insights for more effective and sustainable vector control. Further investigations on mosquito vectorial competence would give more precise insights into the impacts of this vector control strategy on malaria transmission.

## Acknowledgements

We extend our sincere gratitude to all research team members at the Laboratory of Vector-borne Infectious Diseases, Regional Institute of Public Health/University of Abomey-Calavi, Benin, for their help with the insectarium work. We are also grateful to Priscille Barreaux and Geraldine Foster of Department of Vector Biology, Liverpool School of Tropical Medicine for technical support.

## Author Contributions

**Conceived and designed the experiments**: Pierre Marie Sovegnon, Luc Salako Djogbénou.

**Performed the experiments**: Pierre Marie Sovegnon, Marie Joelle Fanou.

**Data analysis:** Pierre Marie Sovegnon, Oswald Yédjinnavênan Djihinto.

**Writing – original draft:** Pierre Marie Sovegnon. Romaric Akoton.

**Writing – review & editing:** Pierre Marie Sovegnon, Romaric Akoton, Oswald Yédjinnavênan Djihinto, Hamirath Odée Lagnika, Romuald Agonhossou, Luc Salako Djogbénou.

